# Axonal selectivity of myelination by single oligodendrocytes established during development in mouse cerebellar white matter

**DOI:** 10.1101/2023.10.03.560586

**Authors:** Batpurev Battulga, Yasuyuki Osanai, Reiji Yamazaki, Yoshiaki Shinohara, Nobuhiko Ohno

## Abstract

Myelin formation by oligodendrocytes regulates conduction velocity and functional integrity of neuronal axons. While individual oligodendrocytes form myelin sheaths around multiple axons and control the functions of neural circuits in the brains, it remains unclear if oligodendrocytes selectively form myelin sheaths around the specific types of axons. In this study, we developed a method for observing a single oligodendrocyte and its myelin sheaths around different types of axons in the mouse cerebellar white matter. This was achieved by combining sparse fluorescent labeling of oligodendrocytes by attenuated rabies virus and subsequent immunostaining for axonal markers along with tissue clearing. We revealed that approximately half of the oligodendrocytes showed a preferential myelination of axons originating from Purkinje cells in the adult mice. The preference for Purkinje cell axons was more pronounced during development, when the process of myelination within cerebellar white matter was initiated; over 90% of oligodendrocytes preferentially myelinated the Purkinje cell axons. The transgenic mice that label early born oligodendrocytes showed that their myelin sheaths were predominantly formed around Purkinje cell axons in the adult cerebellar white matter. These results suggest that a significant proportion of oligodendrocytes preferentially myelinate functionally distinct axons in the cerebellar white matter, and the axonal preference of myelination is established during development.

## Introduction

Oligodendrocytes in the central nervous system (CNS) form myelin sheaths, which enhance axonal conduction velocities and are critical for brain functions. It has been established that oligodendrocytes play critical roles in maintaining the functional integrity of ensheathed axons (Nave, 2010). Additionally, it has been suggested that myelin ensheathment can affect axonal conduction via oligodendrocyte depolarization (Yamazaki et al., 2019; Yamazaki et al., 2007), and myelin structures critical for regulation of axonal conduction are modulated at the single oligodendrocyte level depending on axonal activities in the microenvironment (Osanai et al., 2018). Given that each oligodendrocyte extends multiple processes and forms myelin sheaths around multiple axons, oligodendrocytes would regulate and are responsible for the functions as well as pathophysiology of the neural circuits in which the ensheathed axons are involved.

Previous studies examined the subtypes of axons myelinated by individual oligodendrocytes. In the cerebral cortex, examination using volume electron microscopy suggested that some oligodendrocytes disproportionately myelinate the axons of inhibitory interneurons, whereas others primarily target excitatory axons or show no bias (Zonouzi et al., 2019). In addition, in the corpus callosum, analyses using volume electron microscopy indicated that individual oligodendrocytes formed compact myelin with similar thickness preferentially around distant axons with a limited range of diameters (Tanaka et al., 2021). In fact, sparse fluorescent labeling of oligodendrocytes in the corpus callosum, along with fluorescent discrimination of axons from different brain regions, showed that some callosal oligodendrocytes preferentially myelinate axons from a particular brain region (Osanai et al., 2017). These previous studies support the concept that individual oligodendrocytes preferentially myelinate specific types of axons, which can be discriminated by structural properties and/or axonal functions (Yang, Michel, Jokhi, Nedivi, & Arlotta, 2020). However, it remains unclear if preferential myelination toward specific axons by oligodendrocyte myelination is a general phenomenon, which types of axons can be preferentially myelinated by oligodendrocytes, and how such preference is generated in the white matter of the brain.

The cerebellum orchestrates complex motor functions, cognitive processes, and balance coordination (Ito, 2008). Cerebellar white matter involves three types of axons (Voogd & Glickstein, 1998). One of them is inhibitory Purkinje cell axons, which correspond to the output of the cerebellar cortex. Others include excitatory climbing fibers and mossy fibers, which are the two major input pathways toward the cerebellar cortex. The input and output fibers of cerebellar cortex pass through the cerebellar white matter and are myelinated by adulthood (Nguyen et al., 2018). In fact, motor functions associated with cerebellum can be severely affected in demyelination and dysmyelination involving the cerebellar white matter (Uchida et al., 2012; Wilkins, 2017). While disturbances or damage to cerebellar myelination affect the functional integrity of the axons, little is known about the myelin formation by each oligodendrocyte toward these different classes of axons in the cerebellar white matter.

In this study, we aimed to address preference toward specific types of axons by individual oligodendrocytes in cerebellar white matter. To achieve this goal, we developed an approach to visualize individual oligodendrocytes and the types of their myelinating axons by combining sparse fluorescent labeling of individual oligodendrocytes using an attenuated rabies virus vector encoding GFP and immunostaining for axonal markers along with tissue clearing. We classified Purkinje cell axons and other types of axons as either myelinated or non-myelinated by GFP-labeled oligodendrocytes, and statistically investigated if there is any myelination preference toward either class of axons. This approach also enabled evaluation of the selectivity during the early stages of development when myelination was just initiated. In addition, using transgenic mice, we examined if the myelination preference of initially matured oligodendrocytes changes in the course of development. Our results demonstrate that a substantial number of oligodendrocytes in the cerebellar white matter preferentially form myelin around Purkinje cell axons, and this preference is established during early development.

## Results

### Oligodendrocytes preferentially myelinate Purkinje cell axons in the cerebellar white matter of adult mice

To identify the neuronal subtypes myelinated by single oligodendrocytes, we sparsely labeled oligodendrocytes in the cerebellar white matter of 8-week-old mice. We used an attenuated rabies virus encoding GFP (RV-GFP) (Osanai et al., 2017). Subsequently, immunohistochemistry (IHC) was performed in combination with a tissue-clearing method to identify the axonal subtypes (Fig. 1A) (Osanai et al., 2022). We found that RV-GFP sparsely labelled single oligodendrocytes in the cerebellar white matter (Fig. 1B). Immunostaining was performed using anti-calbindin antibody, which labels Purkinje cell axons (Jande, Tolnai, & Lawson, 1981), and anti-neurofilament antibody, which labels all axons. This approach distinguished Purkinje cell axons (calbindin (+) / neurofilament (+); hereafter, calbindin (+) axons) and other axons including mossy fibers and climbing fibers (calbindin (-) / neurofilament (+) axons; hereafter, calbindin (-) axons) (Fig. 1C). In general, the number of calbindin (+) axons was lower than that of calbindin (-) axons (calbindin (+) vs (-): 36.49 ± 9.39% vs 63.51 ± 9.39%) (Fig. 1D). Using a combination of RV-GFP and IHC with tissue-clearing, we observed that RV-GFP-labeled oligodendrocytes extended processes and ensheathed both calbindin (+) and (-) axons (Fig. 1E, F), consistent with the previous study that used axonal labeling with fluorescent protein expression (Osanai et al., 2017). We counted the numbers of calbindin (+) and (-) axons ensheathed by RV-GFP-labeled oligodendrocytes (Fig. 1G), as well as those that passed nearby the labeled oligodendrocytes but were not ensheathed by them (Fig. 1H). We used Fisher’s exact test for a careful evaluation of bias toward calbindin (+) or (-) axons in each oligodendrocyte. We found that 8 out of 17 (about 47%) GFP-labeled oligodendrocytes preferentially myelinated calbindin (+) axons (Fig. 1H), while one oligodendrocyte preferentially myelinated calbindin (-) axons (Fig. 1H). These results indicate substantial number of the oligodendrocytes preferentially myelinate calbindin (+) axons in the cerebellar white matter of adult mice.

**FIGURE 1:**
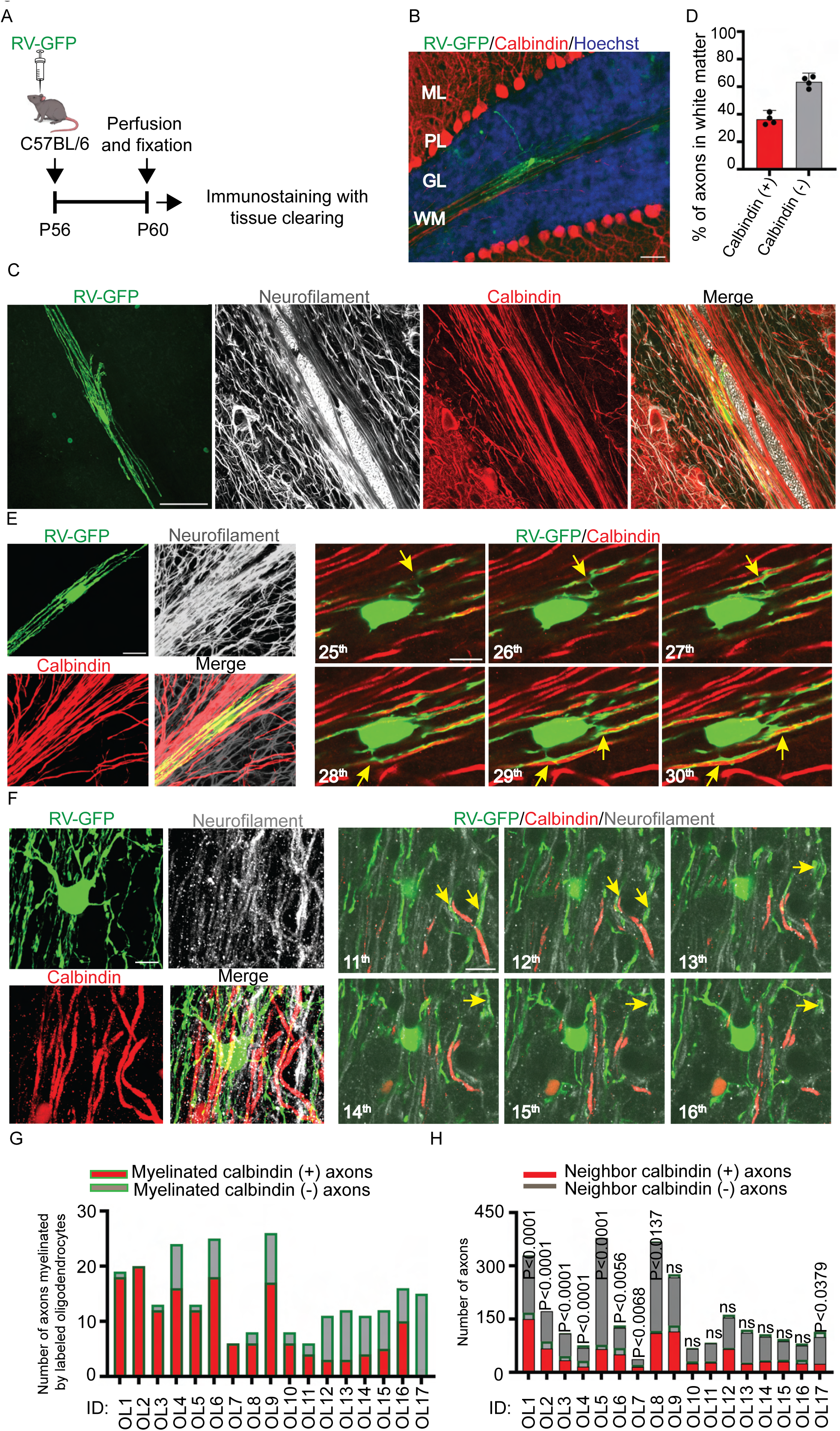
Oligodendrocytes preferentially myelinate Purkinje cell axons in the cerebellar white matter of adult mice. (A) Methods and experimental timeline of injection of rabies virus encoding GFP (RV-GFP) and sample preparation. (B) A representative image showing an oligodendrocyte sparsely labeled with RV-GFP (green) in the cerebellar white matter. Purkinje cells are immunostained for calbindin (red) and nuclei are stained by Hoechst (blue). Scale bar: 50 µm. ML: molecular layer, PL: Purkinje cell layer, GL: granular layer, WM: white matter. (C) A representative image of an RV-GFP-labelled oligodendrocyte (green) and axons immunostained for neurofilament (white) and calbindin (red). Scale bar: 30 µm. (D) The percentage of calbindin-positive / neurofilament-positive (calbindin (+)) and calbindin-negative / neurofilament-positive (calbindin (-)) axons in the cerebellar white matter. (E) The left panel shows Z-stacked confocal microscopy images of calbindin (+) and (-) axons and an RV-GFP-labeled oligodendrocyte which preferentially myelinates calbindin (+) axons. Right panel shows serial confocal microscopy images of the left panel, where the oligodendrocyte ensheaths calbindin (+) axons (yellow arrows). The slice numbers in the image stack are shown at the bottom left corners. Scale bars: 20 µm (left) or 10 µm (right). (F) Left panel shows Z-stacked confocal microscopy images of calbindin (+) and (-) axons and an RV-GFP-labeled oligodendrocyte preferentially myelinating calbindin (-) axons. Right panel shows serial confocal microscopy images of the left panel, where the oligodendrocyte ensheaths calbindin (-) axons (yellow arrows). Scale bars: 20 µm (left) or 10 µm (right). (G) The number of calbindin (+) and calbindin (-) axons myelinated by the single RV-GFP-labeled oligodendrocytes. Each oligodendrocyte is shown using an ID number. Data were obtained from 17 oligodendrocytes from 4 mice. (H) The numbers of calbindin (+) and (-) axons myelinated by each of RV-GFP-labeled oligodendrocyte (Myelinated) and neighbor calbindin (+) and (-) axons not myelinated by the oligodendrocyte (Neighbor). P values of the Fisher’s exact test is shown on the bars. ns: P > 0.05. The same ID numbers represent the same oligodendrocytes (G, H).

Axonal diameter regulates initiation of myelination and a single oligodendrocyte tends to myelinate axons with a discrete range of diameters (Lee et al., 2012; Tanaka et al., 2021). Therefore, we analyzed the diameters of calbindin (+) and (-) axons by measuring the full-width at half maximum (FWHM) of fluorescence intensity (Fig 2A-C) (Call & Bergles, 2021; Osso, Rankin, & Chan, 2021; Stedehouder et al., 2019). We analyzed the diameter of calbindin (+) or (-) axons myelinated or not myelinated by the GFP-labeled oligodendrocytes near each oligodendrocyte. We found that calbindin (+) axons tended to be smaller compared to calbindin (-) axons near each of the oligodendrocytes which dominantly myelinated calbindin (+) axons or myelinated calbindin (+) and (-) axons without significant preference (Fig. 2D). In addition, overall diameters of calbindin (+) axons were smaller than that of calbindin (-) axons regardless of the myelin ensheathment by the GFP-labeled oligodendrocytes (Fig 2E). These results suggest that the preference toward calbindin (+) axons is not attributable to the preferential myelination toward larger diameter axons.

**FIGURE 2:**
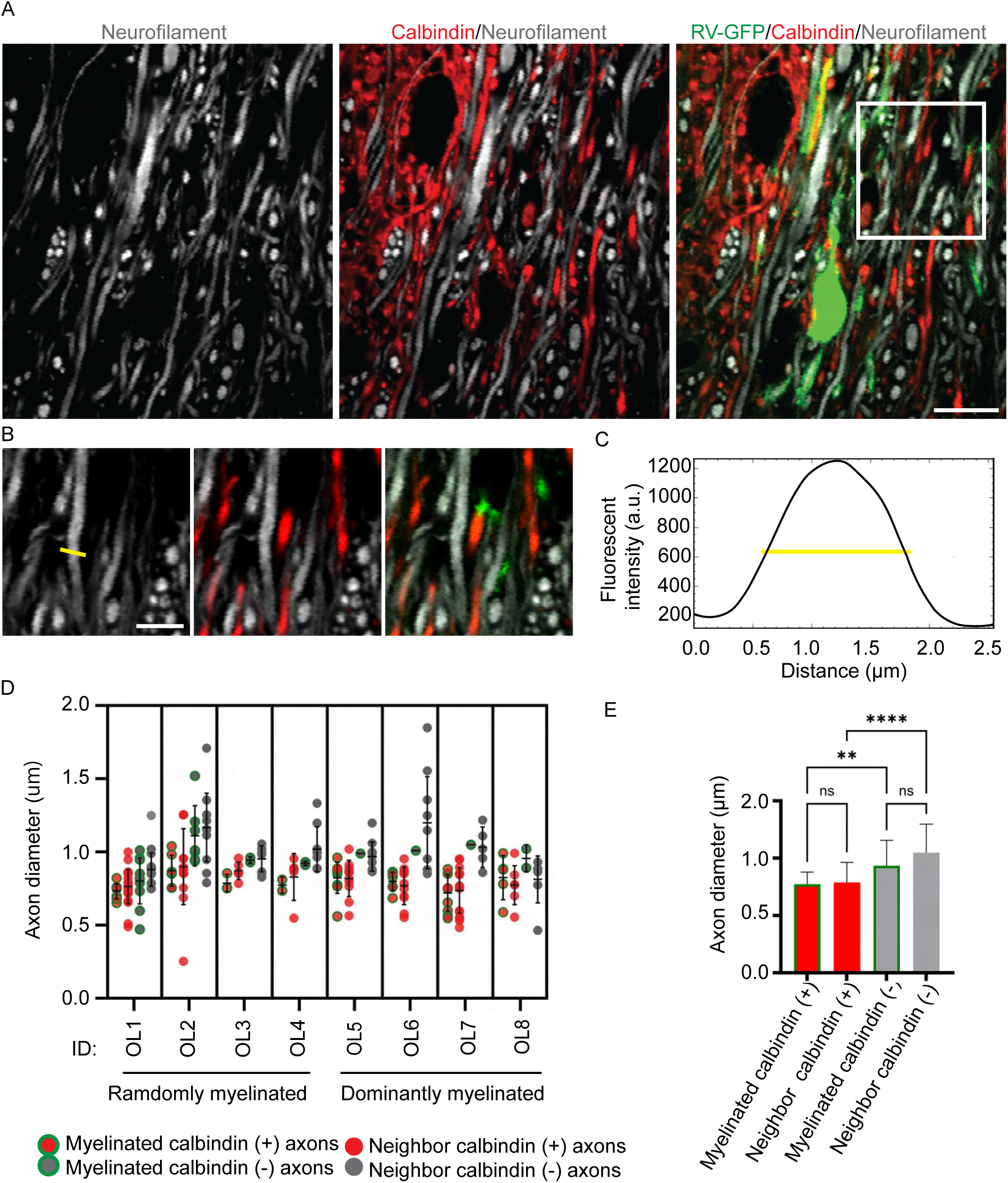
Diameters of axons myelinated and not myelinated by labeled oligodendrocytes are similar. (A) Representative fluorescence images of an oligodendrocyte labeled with rabies virus encoding GFP (RV-GFP) and axons immunostained for calbindin (red) and neurofilament (white). Axonal diameter is measured in the magnified image (B). Scale bar: 15 µm. (B) The magnified image (A, white rectangle) where the diameter analysis is performed using full width at half maximum of fluorescence intensity (yellow line). Scale bar: 5 µm. (C) A fluorescence intensity histogram of the axon indicated in the magnified image (B, yellow line). (D) The diameter of axons myelinated by single oligodendrocytes without any preference to classified axons (OL1-OL4, Randomly myelinated) or with preference to specific axons (OL5-OL8, Dominantly myelinated). Axons are classified into calbindin-positive / neurofilament-positive (calbindin (+)) or calbindin-negative / neurofilament-positive (calbindin (-)) axons, and axons myelinated by each of RV-GFP-labeled oligodendrocyte (Myelinated) or neighbor calbindin (+) and (-) axons not myelinated by the oligodendrocyte (Neighbor). Each dot represents one axon, and bars show the mean ± SD. (E) The diameter of calbindin (+) and (-) axons myelinated and not myelinated by GFP-labeled oligodendrocytes. **: P <0.01, ****: P <0.0001, ns: P > 0.05, one-way ANOVA with Dunn’s multiple comparisons test, N = 35, 65, 20, and 50 for Myelinated calbindin (+), Neighbor calbindin (+), Myelinated calbindin (-), and Neighbor calbindin (-) axons, respectively.

### Oligodendrocytes preferentially myelinate Purkinje cell axons at early developmental period

It is known that myelination of the mouse cerebellum is beginning around P5 to P8 (Chiba et al., 2016; Groteklaes, Bönisch, Eiberger, Christ, & Schilling, 2020). To analyze preference of myelination at the early-stage of cerebellar myelination, RV-GFP was injected into the white matter of P8 mouse cerebellum, and the mice were sacrificed at P12 to identify the axonal subtypes myelinated by the labeled oligodendrocytes with IHC (Fig. 3A). In the cerebellar white matter, GFP-labelled oligodendrocytes were positive for a mature oligodendrocyte marker, CC1, and their processes formed myelin sheaths flanked by a paranodal marker, Contactin-associated protein (Caspr, Fig. 3B). The GFP-positive myelin ensheathed the calbindin (+) and (-) axons, as observed in the adult cerebellar white matter (Fig. 3C). Surprisingly, we found that 22 out of 23 oligodendrocytes preferentially myelinated the calbindin (+) axons (Fig. 3D, E). On the other hand, the diameter of myelinated calbindin (+) and (-) axons were similar at P12 (Fig. 3F). If most of oligodendrocytes predominantly myelinate calbindin (+) axons, it is possible that myelin sheath is predominantly formed on calbindin (+) axons at the beginning of cerebellar myelination. Indeed, when percentages of myelinated calbindin (+) and calbindin (-) axons were analyzed using immunostaining for a myelin marker, myelin basic protein (MBP), we found that 81.34 ± 9.2% of myelinated axons were calbindin (+) axons in the cerebellar white matter at P12 (Fig. 3G, H). Taken together, these data indicate that early born oligodendrocytes selectively myelinate calbindin (+) axons, which are the major axonal subtype initially myelinated in the cerebellar white matter.

**FIGURE 3:**
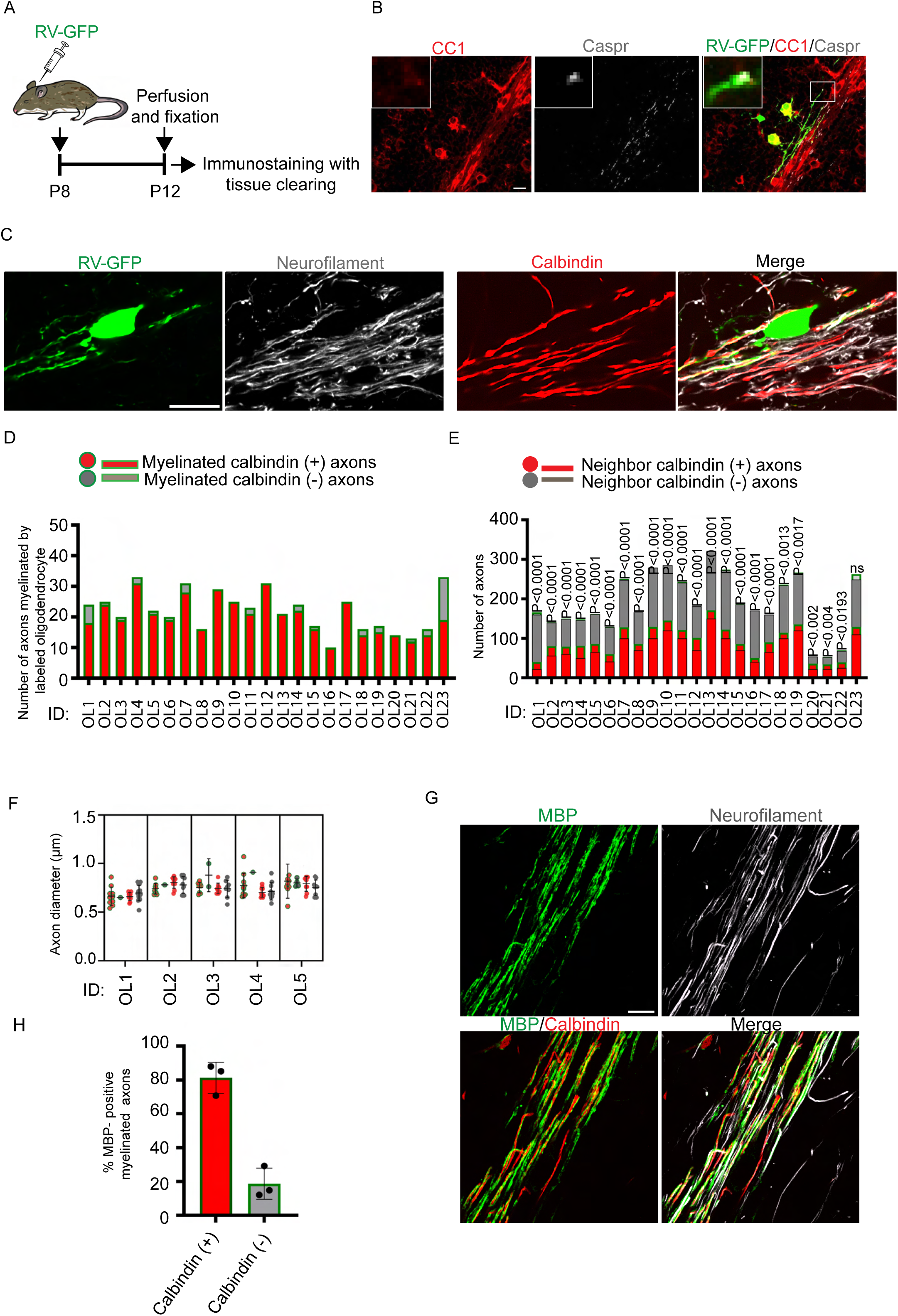
Oligodendrocytes preferentially myelinate Purkinje cell axons at early development. (A) Method and experimental timeline of rabies virus encoding GFP (RV-GFP) injection and sample preparation. (B) GFP-labeled oligodendrocytes were immunopositive for a marker of mature oligodendrocyte, CC1 (red), and the edges of GFP-labelled myelin sheathes were immunopositive for a paranodal marker, contactin-associated protein (Caspr, white). The area marked with a rectangle is magnified in insets. Scale bar: 15 µm. (C) Representative fluorescence images of P12 mouse cerebellar white matter with RV-GFP injection and immunostaining for calbindin and neurofilament. Axons can be classified into calbindin-positive / neurofilament-positive (calbindin (+)) or calbindin-negative / neurofilament-positive (calbindin (-)) axons, and axons myelinated by each of RV-GFP-labeled oligodendrocyte (Myelinated) or neighbor calbindin (+) and (-) axons not myelinated by the oligodendrocyte (Neighbor). Scale bar: 20 µm. (D) The numbers of calbindin (+) and (-) axons which are Myelinated. (E) The numbers of calbindin (+) and (-) axons which are Myelinated or Neighbor. P values of Fisher’s exact test are shown on bars. ns: P>0.05. The data were obtained from 23 oligodendrocytes in 5 mice. (F) Diameter of calbindin (+) and (-) axons which are Myelinated or Neighbor. Each dot represents one axon and bars show mean ± SD. (G) Immunostaining for a myelin marker, myelin basic protein (MBP, green), calbindin (red) and neurofilament (white) in the cerebellar white matter at P12. Scale bar: 20 µm. (H) Percentage of calbindin (+) and (-) axons in total axons ensheathed by MBP immunostaining.

In order to clarify if myelin sheaths were dominantly formed around calbindin (+) axons, we performed serial immunoelectron microscopic analyses in the cerebellar white matter of P12 mice. The fixed cerebellar slices were immunostained for calbindin which was visualized with diaminobenzidine (DAB), embedded in resin following *en bloc* staining, and observed with serial block-face scanning electron microscopy (Fig. 4A). Cerebellar white matter was clearly identified in the immunostained tissue block (Fig. 4B), and in the serial electron microscopic images (Fig. 4C), myelin sheaths were clearly identified around some axons (Fig. 4D). Since treatments which destroy tissue structures and increase antibody penetration were minimized, the DAB signal was present only near the surface of the tissues (Katoh et al., 2017). Therefore, DAB signals were identified near the surface of the tissues, and in the serial images, we tracked the profiles of myelinated axons which were positive or negative for the DAB signals (Fig. 4E). Using this approach, the myelinated axons could be classified into calbindin (+) and (-) in the serial images. Consequently, we found that majority of the myelinated axons in the cerebellar white matter were calbindin (+) at this age (Fig. 4F). Consistent with the immunofluorescence results (Fig. 3H), 83.19 ± 5.3% of myelinated axons were calbindin (+) axons (Fig. 4G). Taken together, these data indicate that oligodendrocytes matured before P12 preferentially myelinate calbindin (+) axons.

**FIGURE 4:**
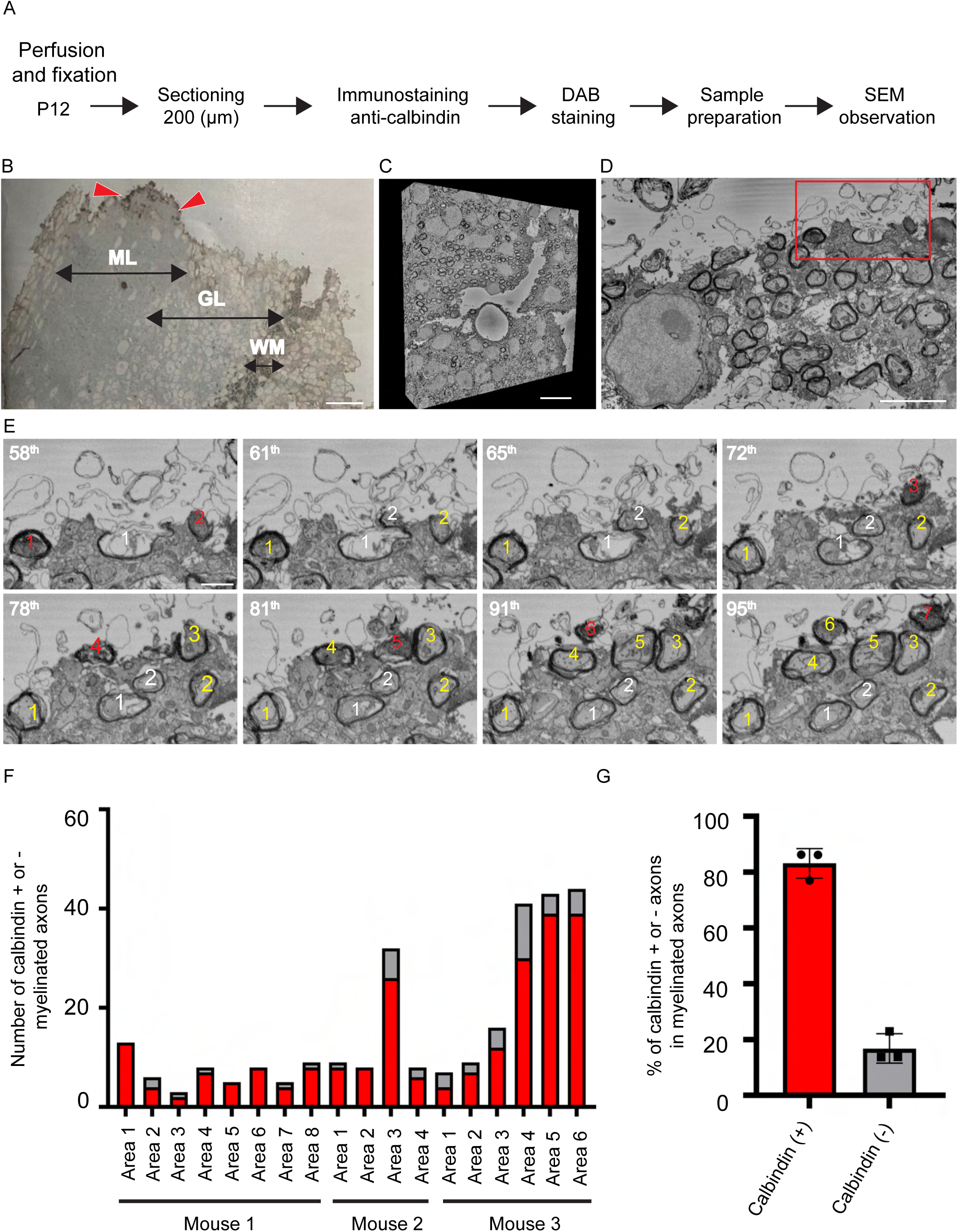
Predominant compact myelin formation around Purkinje cell axons in the cerebellar white matter of P12 mice. (A) An experimental protocol of the serial immunoelectron microscopy. (B) A light microscopic image of a tissue slice immunostained for calbindin with diaminobenzidine (DAB) visualization and embedded in epoxy resin. Myelinated axons are observed in cerebellar white matter (WM) and only the surface of the tissues has DAB signals (arrowheads). ML: molecular layer, GL: granular layer. Scale bar: 20 µm. (C) Three-dimensional reconstruction of the serial immunoelectron microscopic images acquired from the area including cerebellar white matter. Scale bar: 10 µm. (D) One of the serial images of cerebellar white matter including the area indicated with a rectangle and magnified (E). Scale bar: 2 µm. (E) Serial images at a higher magnification showing DAB signals which corresponds to calbindin immunoreactivity in myelinated axons near the surface of the tissue. The slice numbers in the image stack are shown at the upper left corners. Myelinated calbindin-positive (calbindin (+)) and -negative (calbindin (-)) axons are numbered with yellow and white numbers, respectively. The axonal numbers are colored red when DAB signals can be detected in the image. The calbindin (+) axons show DAB-signals in the images near the surface of the tissue but the DAB signal of the profile fade in the deeper areas of the tissue. Scale bar: 500 nm. (F) Numbers of myelinated calbindin (+) and calbindin (-) axons in each of the observed areas from different mice. (G) Percentage of myelinated calbindin (+) and calbindin (-) axons. Each dot represents one mouse.

### Early differentiated oligodendrocytes remain to preferentially myelinate Purkinje cell axons in adult mouse cerebellum

Oligodendrocytes can dynamically form new myelin sheaths in addition to the initially formed sheaths (Duncan et al., 2018; Jeffries et al., 2016; Mezydlo et al., 2023). To address if the preferential myelination of early differentiated oligodendrocytes toward calbindin (+) axons remain in the adult cerebellar white matter, we used *PLP-CreERT/Tau-mGFP* transgenic mice. In these mice, membrane-targeted GFP is expressed only in the mature oligodendrocytes with PLP expression at the timing of tamoxifen exposure, while those matured afterwards are not labeled with GFP (Fig. 5A). In order to label the early born oligodendrocytes, tamoxifen was administrated at P8 and the mice were sacrificed at P56. In these analyses, individual GFP-labeled oligodendrocytes could not be identified because the oligodendrocytes were labeled too densely. However, axonal subtypes ensheathed by GFP-positive myelin could be clearly identified and distinguished from those not ensheathed by the GFP-positive myelin (Fig. 5B). We found that GFP-labeled myelin sheaths were dominantly formed on calbindin (+) axons even in the P56 mouse cerebella (Fig. 5C). In the axons ensheathed by the GFP-labeled myelin sheaths in each mouse, the numbers of calbindin (+) axons exceeded those of calbindin (-) axons (Fig. 5D). However, the number of nearby calbindin (+) axons did not exceed calbindin (-) axons, and the results of Fisher’s exact test for analyzing bias of myelination toward calbindin (+) or calbindin (-) axons showed that GFP-labeled myelin sheaths dominantly formed on the calbindin (+) axons in each mouse (Fig. 5E). These results indicate that early born oligodendrocytes preferentially myelinate calbindin (+) axons in the cerebellar white matter of adult mice.

**FIGURE 5:**
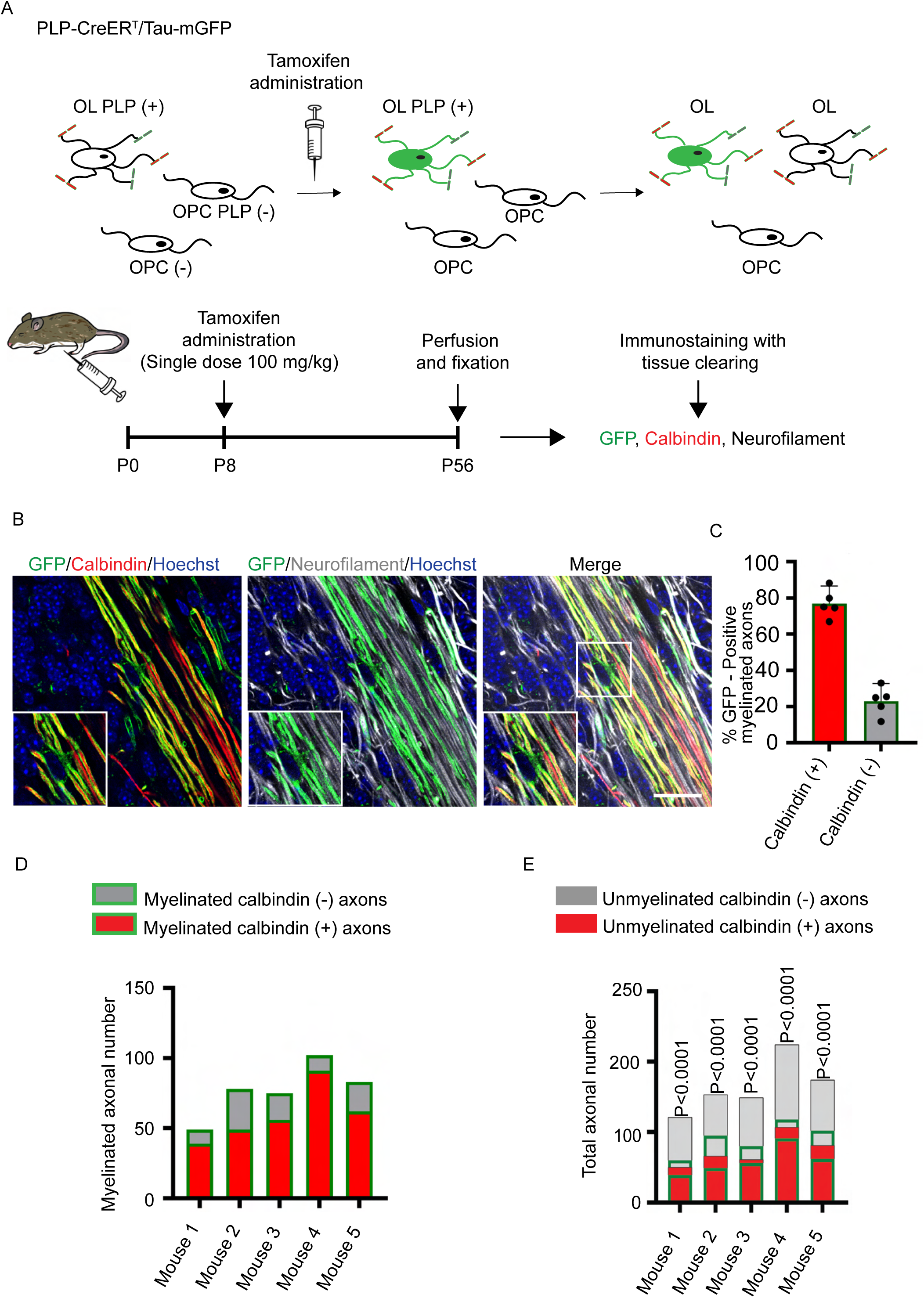
Early differentiated oligodendrocytes remain to preferentially myelinate calbindin (+) axons in adult mouse cerebellum. (A) Schematic diagram of the experimental design using PLP-CreER^T^ mice. OL: oligodendrocyte, OPC: oligodendrocyte precursor cell, PLP: proteolipid protein. In PLP-CreER^T^ mice, the OL differentiated upon tamoxifen injection are labeled with GFP, while those differentiated afterwards are not labeled. (B) Representative immunofluorescence images of a PLP-CreER^T^ transgenic mouse brain section. Myelin sheaths of GFP-labeled oligodendrocytes (green) cover axons double-positive for calbindin (red) and neurofilament (white) (calbindin (+)) and axons positive for neurofilament but not for calbindin (calbindin (-)). Scale bar: 20 µm. (C) The percentage of calbindin (+) and (-) axons ensheathed by GFP-labeled myelin in the cerebellar white matter (76.93 ± 7.82%, 23.07 ± 7.82%, respectively). Each dot represents one mouse. (D) The numbers of calbindin (+) and (-) axons ensheathed by GFP-labeled myelin in each mouse. (E) The numbers of calbindin (+) and (-) axons which are ensheathed or not ensheathed by GFP-labeled myelin in each mouse. P values in Fisher’s exact test are shown on bars.

## Discussion

In this study, we combined labeling of single oligodendrocytes with an attenuated rabies virus vector and immunostaining with a tissue clearing technique, and found that a certain population of oligodendrocytes preferentially myelinate Purkinje cell axons in the cerebellar white matter. We also showed that those oligodendrocytes with preference are differentiated at early developmental stage (< P12). To the best of our knowledge, this is the first report to demonstrate axonal selectivity in myelination by individual oligodendrocytes in the cerebellum and explain how selective myelination observed in the adulthood is established.

Combination of sparse oligodendrocyte labeling and immunohistochemical classification of specific axons enabled thorough examination of axonal preferences of each oligodendrocytes and stringent statistical evaluation. Structural analyses of individual oligodendrocytes including their myelin sheaths has been difficult in the white matter, where many axons and oligodendrocytes are densely distributed. However, sparse labeling of oligodendrocytes has made it possible to observe the detailed morphology of individual oligodendrocytes. Labeling of axons from different brain regions with viral vectors expressing different fluorescent proteins revealed that the proportion of the differentially labeled axons myelinated by individual oligodendrocytes (Osanai et al., 2017). This method appeared applicable to many brain regions even when axonal markers for immunohistochemistry were not available. However, efficiency of axonal labeling with AAV is low, and statistical comparisons of the preference to specific axons are difficult, since the proportion of unlabeled axons is unknown. By contrast, the immunohistochemical labeling visualizes most axons, and therefore it is possible to classify axons with specific markers and identify most axons myelinated or not myelinated by the labeled oligodendrocytes. In addition, the rigorous statistical evaluation can provide reliable results regarding axon preference. On the other hand, a major limitation of this study that using a transgenic mouse to label oligodendrocytes differentiated at the specific developmental stage is that it was difficult to isolate and identify individual oligodendrocytes and sheathes when the density of labeled oligodendrocytes is too high. Refined labeling strategies and also improved tools to visualize more diverse subtypes of axons would provide detailed maps of axonal preference in myelination by individual oligodendrocytes

The clear preference toward Purkinje cell axons was observed in a substantial number of oligodendrocytes, while preference toward other types of axons was not obvious in the cerebellar white matter. Previous studies reported that oligodendrocytes can wrap myelin around cylindrical or fibrous materials, proposing the concept that oligodendrocytes are unselective and use physical cues for myelination (Almeida, 2018; Lee et al., 2012). However, myelination must be finely adjusted for axons within neuronal networks, including elaborate spatial organization and structural modulation of myelin to precisely control arrival timings of action potentials (Almeida & Lyons, 2017; Richardson, McIntyre, & Grill, 2000; Yamazaki et al., 2019; Yamazaki et al., 2014). In fact, recent studies have suggested that cell adhesion molecules, expressed in myelin-forming cells and neurons, can specifically interact with one another and facilitate or repel myelination. (Djannatian et al., 2019; Elazar et al., 2019; Sukhanov, Vainshtein, Eshed-Eisenbach, & Peles, 2021). It is possible that preference toward particular types of axons is a general phenomenon and mediated by mechanisms involving attractive or repulsive guidance of selective myelin formation.

The preference toward Purkinje cell axons was more prominent immediately after the initiation of myelination. The findings that early-born oligodendrocytes preferentially myelinate a particular subtype of neuronal axons is consistent with the developing zebrafish spinal cord, where Mauthner axons are first to be myelinated (Almeida, Czopka, Ffrench-Constant, & Lyons, 2011). It is possible that the axonal subtypes such as Mauthner axons and Purkinje cell axons that are first to be myelinated have distinct functions in the white matter. Purkinje cells can be divided into two groups which express or don’t express aldolase C (zebrin II), and the axons of these cells have different electrical properties and exhibit differential functions (Brochu, Maler, & Hawkes, 1990; Zhou et al., 2014). The outputs through these functionally different axons may require differential regulation of axonal conduction, and important target of selective myelination. In addition, the maintenance of preferential myelination by early differentiating oligodendrocytes until adulthood raises the hypothesis that the preference observed in the adulthood is regulated by the timing of myelination during development; early myelinating and late myelinating oligodendrocytes may determine the neural circuits regulate through preferential myelination. If so, any developmental disturbance and/or event affecting myelination would affect the normal patterns of axonal selectivity and modulate the roles of oligodendrocyte myelination in the conduction regulation and the group of axons affected by oligodendrocyte abnormality in the adulthood. Further studies are required for revealing functional consequence of the preferential myelination toward a particular class of axon.

## Materials and Methods

### Animals and tamoxifen injection

Female C57BL/6 mice and C57BL/6 mouse pups at postnatal days 12 and 19 (P12, P19) were purchased from Japan SLC (Shizuoka, Japan). PLP-CreER^T^ mice (Doerflinger, Macklin, & Popko, 2003) and Tau^mGFP^ mice (Hippenmeyer et al., 2005) were obtained from the Jackson Laboratory (JAX stock # 005975 and # 021162, respectively). PLP-CreER^T^ transgenic mice were crossbred with Tau^mGFP^ homozygous mice to generate PLP-CreER^T^ (either homozygous or heterozygous) and Tau^mGFP^ (heterozygous) mice. Tamoxifen (Nacalai tesque, Kyoto, Japan) was dissolved in corn oil (Sigma, St. Louis, USA) to a final concentration of 10 mg/ml. PLP-CreER^T^ : Tau^mGFP^ mice were injected intraperitoneally with 100 mg/kg tamoxifen at P8. All experiments were approved by the Institutional Animal Care and Use Committee (IACUC) and conducted in accordance with the guidelines for the care and use of animals at Jichi Medical University.

### Injection of RV-GFP into the cerebellum

For visualizing individual oligodendrocytes, we used RV-GFP (Mori & Morimoto, 2014). The procedures of RV-GFP injection have been described previously in detail (Osanai et al., 2017; Osanai et al., 2018). Briefly, 8-week-old C57BL/6 mice were anesthetized with an intraperitoneal injection of mixed anesthetic (0.3 mg/kg medetomidine, 4.0 mg/kg midazolam, and 0.5 mg/kg butorphanol) and placed in a stereotaxic frame (Narishige, Tokyo, Japan). A solution of RV-GFP (1 µl, 3.3 x 10^4^ IU) was stereotaxically injected into lobules IV/V of the cerebella (6.45 mm posterior and 1.00 mm lateral to the bregma, a depth of 0.5 mm). For P8 pups, anesthesia was induced using 3% isoflurane in a plastic chamber, followed by maintenance with 2.5% isoflurane during the stereotaxic surgery. RV-GFP (0.5 µl, 1.7 x 10^4^ IU) was stereotaxically injected into the P8 cerebella (2.7 mm posterior and 0.44 mm lateral to the lambda, a depth of 0.35 mm), using pulled glass pipettes with an inner diameter of 20-40 µm and an air pressure system. The injection took about 3 minutes. The mice were sacrificed 4 days after the RV-GFP injection.

### Immunohistochemistry

Mice were perfused with 4% paraformaldehyde in 0.1M phosphate buffer (PB) at P56 for PLP-CreER^T^:Tau^mGFP^ transgenic mice and at P12 and P56 for C57BL/6 mice. Brains were embedded in 5% agarose in phosphate-buffered saline (PBS) prior to slicing, and a vibratome slicer (LinearSlicer PRO 10; DSK, Kyoto, Japan) was used to obtain 70 µm-thick brain slices. Lobules IV/V of the cerebella were carefully dissected from the brain slices using a liner paintbrush. The cerebellar slices were then postfixed in the same fixative for 24 hours at 4°C. The slices were washed three times in PBS for 10 minutes each at room temperature (RT), and immunostaining was performed according to the manufacturer’s protocol of RapiClear 1.52 (RC152001, Sunjin Laboratory, Taiwan, China) with minor modifications. The slices were incubated in 2% PBST (2% Triton X-100 and 0.05% sodium azide in PBS) for 24 hours at RT. They were then washed three times with PBS for 10 minutes each and incubated in blocking buffer (10% normal goat serum, 1% Triton X-100, 2.5% DMSO and 0.1% sodium azide in PBS) for 24 hours at RT. The slices were further incubated with various antibodies, including chicken anti-neurofilament-H (1/500, NB300-217, Novusbio, Colorado, USA), rabbit anti-calbindin (1/500, MSFR100390, Frontier Institute, Hokkaido, Japan), anti-GFP (Rat IgG2a) (1/500, 04404-84, Nacalai tesque), anti-APC (CC1, 1/100, OP80, Merk, Darmstadt, Germany) and rat anti-MBP (1/100, MCA409S, Bio-Rad, California, USA) in antibody dilution buffer (1% normal goat serum, 0.2% Triton X-100, 2.5% DMSO and 0.1% sodium azide in PBS) for 3-4 days at 4°C. The anti-GFP antibody was used to visualize GFP in the Tau^mGFP^ mice. They were then washed three times with washing buffer (3% NaCl and 0.2% Triton X-100 in PBS) for 1 hour each at RT. The cerebellar slices were subsequently incubated with secondary antibodies in antibody dilution buffer, including goat anti-chicken IgY Alexa Fluor 647 (1/200, A-21449, ThermoFisher, Massachusetts, USA), goat anti-rat IgG Alexa Flour 488 (1/200, A-11006, ThermoFisher) and goat anti-rabbit IgG Alexa Flour 568 (1/200, A-11011, ThermoFisher), for 2 days at 4°C. They were then washed three times with washing buffer for 1 hour each. The cerebellum slices were incubated with Hoechst 33342 (1/2000, H3570, ThermoFisher) for 2 h at RT. After a final wash with PBS for 10 minutes, the slices were placed on glass slides and each slice was treated with 15-20 µl of RapiClear solution. Finally, the slices were mounted with RapiClear solution.

### Confocal microscopic analysis

Series of Z-stack projections were acquired at 0.5 µm intervals (objective lens ×60 or ×100; N.A., 1.42 or 1.45, respectively) using Dragonfly high speed confocal microscope system (Oxford instruments, Abingdon-on-Thames, UK) or FV1000 (objective lens ×60; N.A., 1.35) (Olympus, Tokyo, Japan). Analysis of individual oligodendrocytes and counting of axonal numbers were done using serial confocal microscopy images. We identified the top and bottom parts of each oligodendrocyte by detecting the disappearance of the RV-GFP signal. We also determined the outermost processes using stacked serial images. Axons adjacent to the oligodendrocytes were defined as those falling within the range between the outermost processes, as calculated from the stacked serial confocal images. RV-GFP (+) oligodendrocyte processes that are wrapping around axons labelled with anti-calbindin and/or anti-neurofilament antibodies were defined as myelin sheaths. The diameter of axons was calculated by measuring the full-width at half maximum (FWHM) of fluorescence intensity using Fiji software. Briefly, five lines perpendicular to a targeted axon were drawn using Fiji, and histograms of fluorescence intensity for each line were made using the Plot profile function of Fiji. From the Gaussian fit of fluorescence intensity histogram, FWHM was calculated for each line. The average of FWHM of the five lines was calculated and determined as the diameter of targeted axon.

### Immunoelectron microscopy analysis

P12 C57BL/6 mice were anesthetized and perfused transcardially with saline, followed by a 4% paraformaldehyde and 2.5% glutaraldehyde solution in PB. The brains were removed and postfixed overnight in the same fixative. Then, they were cut into 200-µm-thick sagittal slices using a vibratome slicer. Slices of the targeted region in the cerebellum (lobules 4/5) were gently dissected from the brain sections using a razor blade. The slices were washed with PBS four times for 2 minutes each at 4 °C, and then incubated in 5% bovine serum albumin (BSA) in PBS for 3 hours at room temperature (RT). Subsequently, the cerebellar slices were incubated in rabbit anti-calbindin (1/50, MSFR100390, Frontier Institute) diluted in PBS containing 5% BSA for 3 hours at RT. The slices were then washed with PBS three times for 3 minutes each at RT. Next, the cerebellar slices were incubated in an HRP-conjugated anti-rabbit IgG (1:100, 111-035-144; Jackson ImmunoResearch, Pennsylvania, USA) for 1 hour at RT and washed with PBS three times for 3 minutes each at RT. Finally, the cerebellar slices were incubated in DAB solution (0.5 mg/ml DAB with 10 nM H_2_O_2_ in PBS) for 10 minutes, and then washed with PBS three times for 3 minutes each at RT. Thereafter, the samples were stained *en bloc*, dehydrated and embedded in resin, as described previously (Morizawa et al., 2022). Slices were washed 4 times in ice cold PBS and then treated with PBS containing 2% OsO_4_ (Nisshin EM, Tokyo, Japan) and 1.5% K[Fe(CN)_6_] (Nacalai tesque) on ice. The slices were then incubated with 1% thiocarbohydrazide (Tokyo Chemical Industry, Tokyo, Japan) for 20 minutes and then with 2% OsO_4_ for 30 minutes at RT. The slices were treated with 2% uranyl acetate at 4°C overnight and for an additional 30 minutes at RT. They were then stained with Walton’s lead aspartate at 50°C for two hours. Each of these treatments was followed by washing four times in water. The slices were then dehydrated with a graded ethanol series (60%-80%-90%-95%) at 4°C, infiltrated sequentially with dehydrated acetone, a 1:1 mixture of acetone and Durcupan (Sigma), a 1:3 mixture of acetone and Durcupan, and finally 100% Durcupan. The incubation in Durcupan was done overnight and then repeated in fresh Durcupan for an additional 90 minutes. Then Durcupan was polymerized at 60°C for 3 days. The pieces of Durcupan with samples were attached on aluminum rivets, trimmed and imaged with a field emission type scanning electron microscope equipped with the 3View system and the OnPoint backscattered electron detector (Gatan, Pleasanton, CA). Serial images were acquired at the resolution of 5.0 nm/pixel and 50 nm steps in the depth direction. The images were processed using Fiji (https://fiji.sc/) and Amira (ThermoFisher).

### Statistical Analysis

Statistical analyses were performed using Prism 7 (GraphPad Software). Using confocal images with GFP-labeled oligodendrocytes and their processes and myelin sheaths, calbindin (+) and (-) axons were identified, and the numbers of calbindin (+) and (-) axons ensheathed and not ensheathed by GFP-labeled myelin were counted. Using the numbers, Fisher’s exact test was performed for accessing preference toward calbindin (+) or (-) axons. One-way ANOVA with Dunn’s multiple comparisons was used for nonparametric multiple comparison. Boxes and bars in the graphs show mean ± SD.

## Acknowledgement

We thank Dr. Shinsuke Koyama (The Institute of Statistical Mathematics) for advice of statistical analysis. We thank Mr. Tom Kouki and Ms. Megumi Yatabe (Jichi Medical University) for their technical assistance. This study was supported by KAKENHI Grants from the Japan Society for the Promotion of Science, (21H05241, 21H04786 and 20KK0170) to NO, (21K15197) to YO, and a research grant from the National Center of Neurology and Psychiatry (No. 3–5) to NO., a research grant from The Uehara Memorial Foundation (202210148) to YO.

